# Mapping the content of comments on bioRxiv and medRxiv preprints

**DOI:** 10.1101/2022.11.23.517621

**Authors:** Clarissa F. D. Carneiro, Gabriel Costa, Kleber Neves, Mariana B. Abreu, Pedro B. Tan, Danielle Rayêe, Flávia Boos, Roberta Andrejew, Tiago Lubiana, Mario Malički, Olavo B. Amaral

**Affiliations:** Institute of Medical Biochemistry Leopoldo de Meis, Universidade Federal do Rio de Janeiro, Rio de Janeiro, Brazil; BIH QUEST Center for Transforming Biomedical Research, Berlin Institute of Health, Germany; Carlos Chagas Filho Institute of Biophysics, Universidade Federal do Rio de Janeiro, Rio de Janeiro, Brazil; Department of Ophthalmology and Visual Sciences, Albert Einstein College of Medicine, NY, USA; Programa de pós-graduação em Psicobiologia, Universidade Federal de São Paulo, Brazil; Department of Biochemistry, Institute of Chemistry, Universidade de São Paulo, São Paulo, Brazil; Ronin Institute, virtual organization; School of Pharmaceutical Sciences, University of São Paulo, São Paulo, Brazil; Meta-Research Innovation Center at Stanford (METRICS), Stanford University, Stanford, CA, USA

**Keywords:** Preprints, preprint feedback, preprint review, peer review, post-publication peer review

## Abstract

**Introduction:** Preprints have been increasingly used in biomedical sciences, providing the opportunity for research to be publicly assessed before journal publication. With the increase in attention over preprints during the COVID-19 pandemic, we decided to assess the content of comments left on preprint platforms.

**Methods:** Preprints posted on bioRxiv and medRxiv in 2020 were accessed through each platform’s API, and a random sample of preprints that had received between 1 and 20 comments was analyzed. Comments were evaluated in triplicate by independent evaluators using an instrument that assessed their features and general content.

**Results:** 7.3% of preprints received at least 1 comment during a mean follow-up of 7.5 months. Analyzed comments had a median size of 43 words. Criticisms, corrections or suggestions were the most prevalent type of content, followed by compliments or positive appraisals and questions. Most critical comments regarded interpretation, data collection and methodological design, while compliments were usually about relevance and implications.

**Conclusions:** Only a small percentage of preprints posted in 2020 in bioRxiv and medRxiv received comments in these platforms. When present, however, these comments address content that is similar to that analyzed by traditional peer review. A more precise taxonomy of peer review functions would be desirable to describe whether post-publication peer review fulfills these roles.

## Introduction

Preprints have been proposed as a way to accelerate scientific communication for many years (Cobb, 2017); however, only recently has the format gained attention from the biomedical science community (Berg *et al*., 2016). Their adoption in this field has grown since the 2010s (*Biology preprints over time*, 2020), and increased sharply after the COVID-19 pandemic (Fraser *et al*., 2021).

A limitation of preprints, which has been the target of much criticism, is the lack of peer review (Sheldon, 2018; Puebla, Polka and Rieger, 2022; Sever, 2022). Nevertheless, accumulating evidence suggests that the quality of preprints is not markedly inferior compared to peer-reviewed journal publications, and that changes between preprints and published versions are usually minor (Carneiro *et al*., 2020; Bero *et al*., 2021; Akbaritabar, Stephen and Squazzoni, 2022; Brierley *et al*., 2022).

On the other hand, a frequently mentioned advantage of preprints is the possibility of rapidly receiving feedback from the community (Sever *et al*., 2019). Nevertheless, evidence for the fulfillment of this role is scarce or anecdotal (Sever *et al*., 2019; Oransky and Marcus, 2020; Puebla, Polka and Rieger, 2022), as community engagement with post-publication peer review platforms is irregular and usually underwhelming (Dolgin, 2018).

An analysis of bioRxiv preprints posted between 2015 and 2019 (Malički *et al*., 2021) estimated that less than 10% of them received any comments. After the COVID-19 crisis, however, an increased attention was directed at preprints, and Fraser et al. reported that 16% of a 2020 sample of COVID-related preprints posted to bioRxiv and medRxiv received comments (Fraser *et al*., 2021). Nevertheless, the specific content of these comments has not been studied in detail.

In the present study, we aim to describe the content of comments received by preprints posted in 2020 to bioRxiv and medRxiv. This study extends upon previous ones by building a predefined content taxonomy based on the findings of previous qualitative studies, in order to provide a more quantitative analysis. This description can be useful in informing future research on preprints and post-publication peer review.

## Methods

A protocol for this study was developed prior to its execution and is available at https://osf.io/54xwy. Deviations from the protocol are described in this section and summarized at https://osf.io/b6up2. All relevant scripts and datasets used are available at https://osf.io/k9e8c/. Analyses were done with R version 4.2.2 (R Core Team, 2022).

### Study sample

All preprints posted to bioRxiv and medRxiv in 2020 were accessed through each platform’s API on March 29^th^, 2021. After filtering those that received at least 1 comment to their first version, we obtained a list with 1,903 preprints from bioRxiv and 1,108 preprints from medRxiv. Preprints with more than 20 comments (5 from bioRxiv and 29 from medRxiv) were excluded, and the remaining ones were randomly sampled until the total number of comments reached 1,000 for each platform. We were not able to obtain data from Research Square as planned in the protocol, as information on comments was not available in the beta version of their API.

### Analysis of comments

A data collection form was built based on previous studies about peer review content (Glonti *et al*., 2019) and preprint comments (Malički *et al*., 2021). An instruction manual was made available to all evaluators and contained detailed information on how data should be collected and classified (https://osf.io/rmjz3).

Comments were evaluated by researchers who initially participated in a pilot study to refine the data collection instrument, including a sample of 17 comments from 10 preprints. Once the data collection form and protocol were finalized, all evaluators had access to a training set of 12 comments that had been consensually evaluated by the coordinating team. Consensus answers were visible after evaluators completed their assessments, to allow for resolution of queries and discrepancies. Comments used in these stages were not included in the final sample.

Each sampled preprint was assigned to 3 evaluators, who filled out the data collection form for all comments posted for that preprint until March 29^th^, 2021. Agreement between evaluators was calculated after completion of data collection. We calculated Fleiss kappa coefficients between all evaluators for each question in the form, as well as the overall percentage of agreement for all questions between each pair of evaluators.

**Fig. 1** summarizes the initial process of comment coding. Briefly, evaluators first identified whether each comment was a response to a previous comment, whether it was made by one of the preprint authors and whether they referred to the content of the preprint. Comments that were responses to other comments did not have their content analyzed.

**Figure 1.**
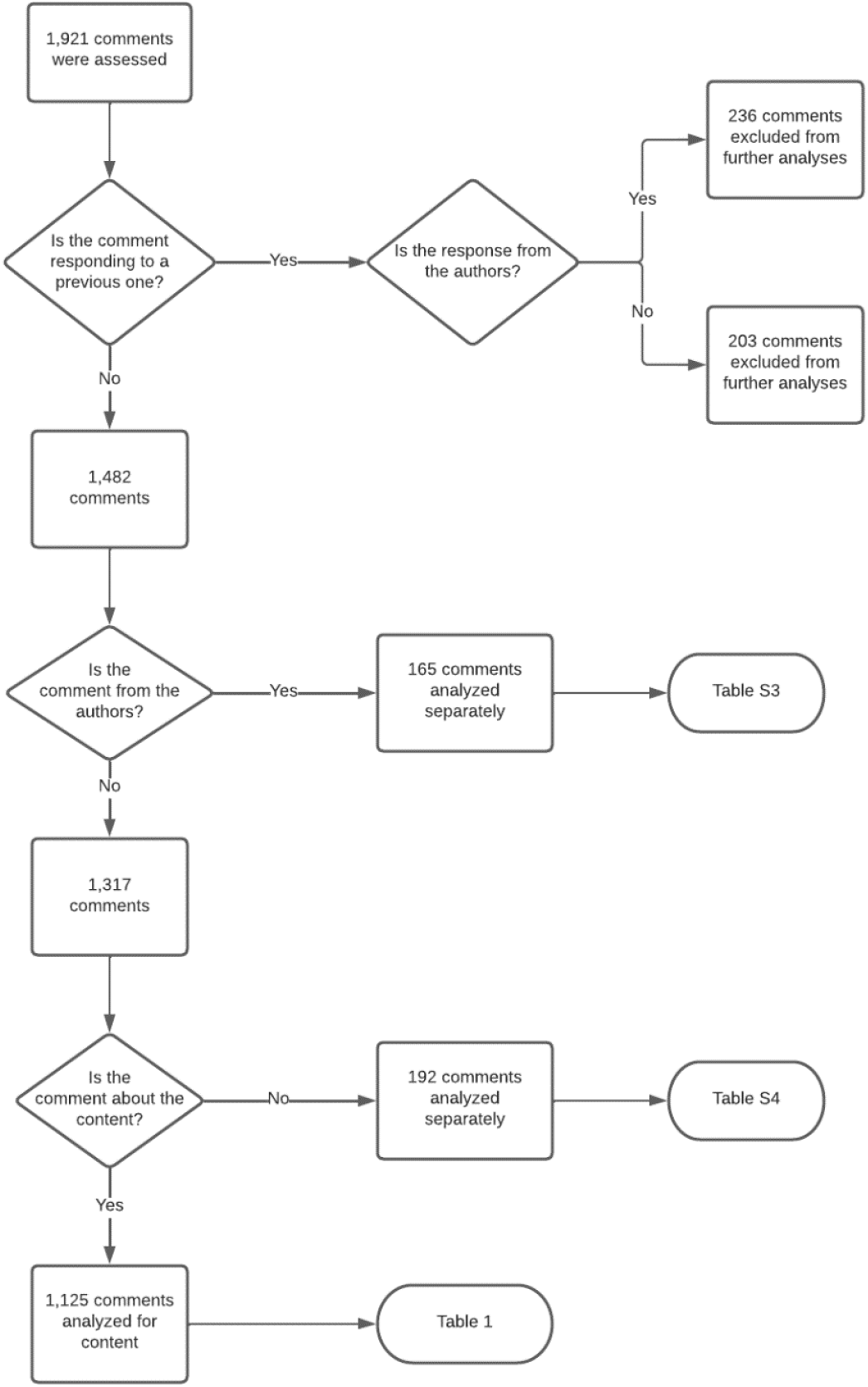
Flow diagram of the analysis process. Responses to other comments were not analyzed. Comments from the preprint authors or that were classified as not being about the preprint content were analyzed using specific categories. All other comments (i.e., not responses, from non-authors, and about the content) were analyzed using the complete data extraction form.

Comments were classified according to their entire content (i.e., classifications were not linked to specific sentences or phrases) using a preset list of categories, and each comment could have several content categories. Comments from the preprint’s authors were classified according to the following predefined categories (adapted from Malički *et al*. 2021): new data, new analyses, additional information, publication status, corrections, feedback request, study promotion and publication venue request. Evaluators also had the option of choosing “other” and briefly describing the content in an open format. After completing data collection, we reviewed these descriptions and created a new category (extra materials). All other descriptions were reclassified under one of the existing categories.

When a comment was classified as not being about the content of the preprint (a category which included comments merely linking to other resources), the evaluator was asked to briefly describe the content of the comment. A classification was derived from these descriptions after completion of data collection, yielding the following categories: link to external resource (peer review, reference, blog, data or media), topic under study, publication status, collaboration proposal, scholarly communication, access to materials, importance of science, public engagement, interaction with stakeholders (media or policymakers), apology, authorship, and conflict of interest.

All other comments (i.e., not responses, not from authors, and about the content) were evaluated using the main data collection form. We initially looked at general format features by asking whether comments were clearly written, whether they included personal offenses (to the authors or in general) and whether they were the product of an organized review effort. We assessed general content by asking whether the comments included (a) criticisms, corrections or suggestions, (b), compliments, (c) questions or (d) other content not fitting these categories. We also asked whether contents contained a summary description, the types of references included (if any), whether they contained new data or analyses and whether they questioned the preprint’s conclusions.

Criticisms, corrections or suggestions, compliments, questions and other general content were further evaluated concerning their specific content. Evaluators were asked to select one or more categories from the following list: Novelty/Originality, Relevance, Readability, Previous literature, Concepts/theoretical framework, Title/abstract, Methodological design, Materials and data collection, Ethics, Data visualization, Analysis, Interpretation, Implications, Data sharing, Reporting, Additional information. They could also select “other” and provide a brief description in an open format. After completion of data collection, we reviewed all descriptions, but no new categories were identified, as most evaluators used the space to elaborate or justify their choices. Comments with no attributed categories were classified under one of the available categories based on these descriptions.

The most prevalent answer for each question with mutually exclusive categories was considered final and used in the analyses. In case of triple disagreements, a fourth evaluator reviewed the questions and made the final decision. This was a deviation from our protocol, as we initially planned to involve all evaluators in discussion, but for simplicity decided on one adjudicator. For questions in which categories were not mutually exclusive - i.e. content of authors’ comments, type of reference and specific content –, agreement was not required, and every option selected by at least one evaluator was included in the analysis.

### Additional data collection

Besides the content collected through the form, we collected the total number of words for each comment, counting hyperlinks as a single word. In addition to the protocol, we also classified the types of organized review efforts identified.

For each preprint, we collected the date of posting, subject area (selected by the authors from each platform’s predefined options), region of origin (defined by the corresponding author’s affiliation), whether it had been withdrawn (data collected between December 1^st^ and 14^th^, 2021) and whether it had been published in a journal (data collected on October 27^th^, 2021). If published, we collected the date, venue of publication and journal’s impact factor (from the 2020 Journal Citation Report). For both preprints and peer-reviewed publications, we collected Altmetric Attention scores and numbers of citations (on October 27^th^ and 28^th^, 2021, respectively). Preprints were also classified as being related to COVID-19 or not by searching for the following keyword in their title or abstract: “coronavirus”, “covid-19”, “sars-cov”, “ncov-2019”, “2019-ncov”, “hcov-19”, “sars-2” (Fraser *et al*., 2021).

### Data analysis

For each question, we present the frequency (number and proportion) of “Yes” answers to general content and format questions among the sampled comments. Proportions are always relative to the total number of comments eligible for assessment in each question, excluding those that were removed in previous data collection steps (**Fig. 1**). For general content and format categories, 95% confidence intervals for these proportions are included. We also present the frequency of “Yes” answers in subsets of comments (comments from organized review efforts, comments that questioned the preprint conclusions and comments that received a response), for a qualitative comparison with the complete dataset, an additional analysis not originally planned in the protocol.

We also include additional analyses of the general content and format categories using the preprint (rather than the comment) as the unit of analysis. For this, we count the number of preprints in which at least one of the comments presented a particular feature in relation to the applicable total of preprints (following the exclusion flow, as described in **Fig. 1**).

To explore associations between comment content and preprint features we built logistic regression models where each content category (taking “No” as the reference) was taken as the response variable and present McFadden’s pseudo-R^2^ values and p values from Analysis of Deviance tests. The sign/direction of effects is presented for quantitative and binomial predictors (using medRxiv for platforms and “Yes” for COVID-related categories as positive reference values). While in the protocol we described that these exploratory associations would be modeled for each question, we restricted our analyses to general questions in which response prevalence was between 5% and 95%. In addition, we restricted the correlation with subject areas to the 10 most frequent areas from each platform.

## Results

### Sample description

We identified 52,736 preprints published in 2020 (38,667 in bioRxiv and 14,069 in medRxiv), 7.3% of which had received at least one comment (2,412 in bioRxiv and 1,440 in medRxiv). Considering only first versions that had received less than 20 comments, there were 3,070 comments for 1,898 bioRxiv preprints and 2,316 comments for 1,079 medRxiv preprints to sample from. We sampled 611 preprints from bioRxiv and 525 preprints from medRxiv to reach a total of 1,000 comments from each platform. Data collection was completed for 1,921 comments from 1,037 preprints, due to discrepancies between the information available on the API and on the preprint webpage. **Fig. 2A** shows the distribution of comments per preprint on each platform. These comments had a median of 43 words, ranging from 1 to 3,172 (**Fig. 2B**).

**Figure 2.**
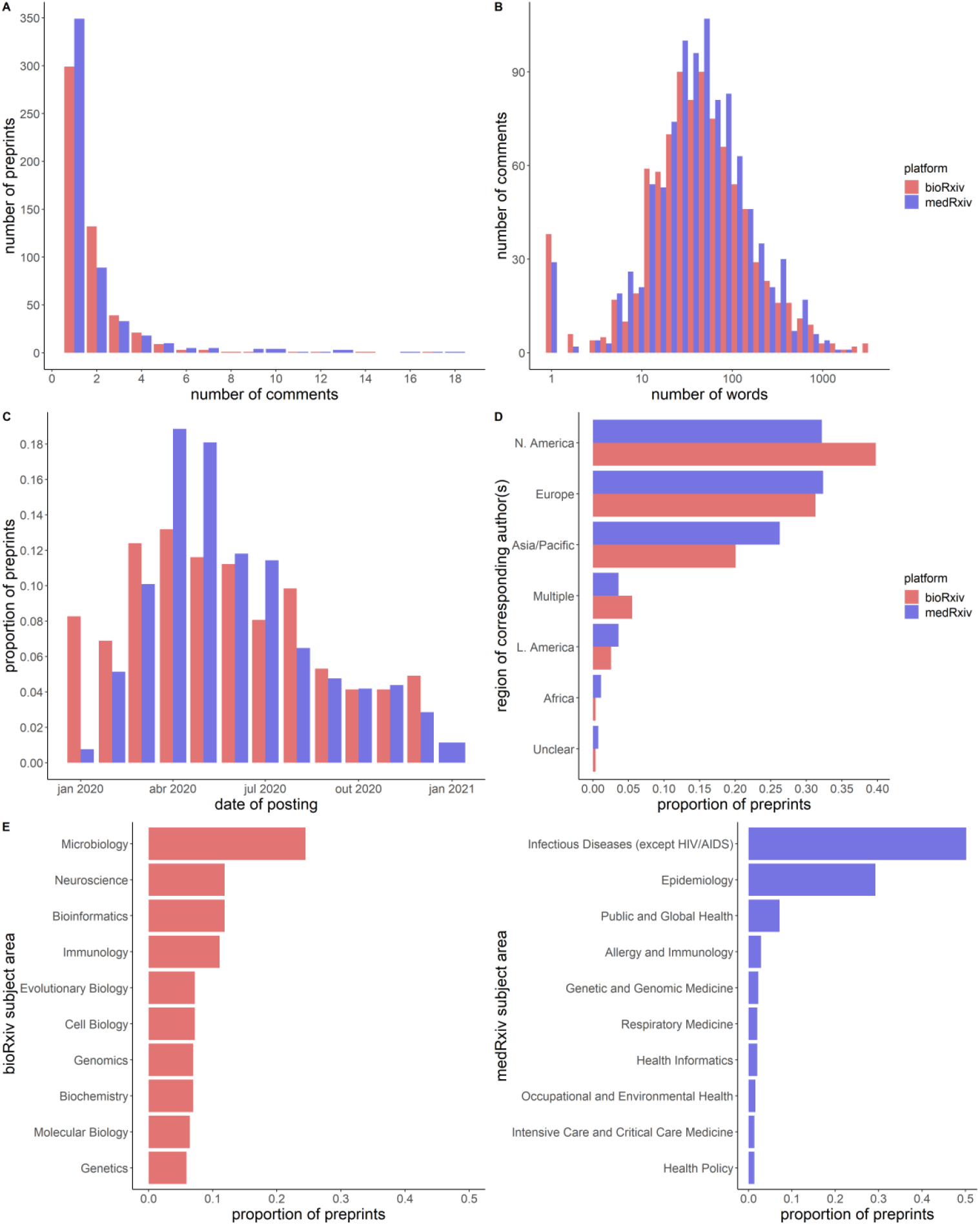
Sample description. Bars represent the total number of preprints in each bin/category, with those from bioRxiv in red and those from medRxiv in blue. **(A)** Distribution of number of comments per preprint in the analyzed sample. The median number of comments is 1 (with a maximum of 17 for bioRxiv and 18 for medRxiv). **(B)** Distribution of comment size in the analyzed sample. The median (range) number of words is 42 (1-3,172) on bioRxiv and 44 (1-1,640) on medRxiv. Overall median is 43 (1-3,172). The peak in 1-word comments mostly consists of those containing isolated hyperlinks. **(C)** Distribution of publication dates, aggregated by month. **(D)** Region of origin of the corresponding authors. ‘Multiple’ combines cases where the corresponding author had affiliations in more than one region or where multiple corresponding authors had affiliations in different regions. Region of origin could not be identified for 18 preprints. **(E)** Areas of research. Given the large number of areas with very few preprints in our sample, we only show the top 10 areas for bioRxiv (*left*) and for medRxiv (*right*).

There is a notable increase in the number of preprints posted between March and June 2020 (**Fig. 2C**), particularly on medRxiv, which coincides with the peak described for COVID-19 preprints across multiple platforms (Fraser *et al*., 2021). Unsurprisingly, most preprints were from authors based in North America and Europe, followed closely by those based in Asia and the Pacific region (**Fig. 2D**). 91% of medRxiv preprints and 35% of bioRxiv preprints in our sample were identified as COVID-related. Because of this, medRxiv concentrated most preprints in infectious diseases and epidemiology, while the most common category in bioRxiv was microbiology. Preprints in our sample had a median of 3 citations (**Suppl. Fig. 1C**), and 23.2% of them did not receive any citations, while their median Altmetric score was 21.2 (**Suppl. Fig. 1B**).

29% of the preprints in our sample had been published in a peer-reviewed journal by October 27^th^, 2021, with a mean of 129 days from preprint to publication. On average, these journals had an impact factor of 11.1. Four preprints in our sample were later withdrawn, but we did not assess whether the comments received had any impact on this decision and did not exclude them from the analyses.

### Content of comments

Agreement between evaluators is presented in **Suppl. Tables 1-2**. Most disagreements occurred when one of the evaluators chose different answers at the initial stages of the form (e.g. whether comments were about the preprint’s content), generating disagreement on subsequent questions as well.

A flowchart describing the evaluation process is provided in **Fig. 1**. Comments identified as responses to other comments (n = 439) were excluded from further analysis. Of the 1,482 remaining comments, 165 (11.1%) were identified as having been posted by one of the preprint’s authors and classified using a separate subset of categories, with detailed results shown in **Suppl. Table 3**. Most commonly, authors used comments to provide information on the preprint’s publication status (54%), i.e. submission, revision, acceptance or publication by a journal, additional information on the study (30%) or corrections to the preprint (18%).

The remaining 1,317 comments were from non-authors. In order to assess their content using a taxonomy suited for peer review, we started by assessing whether the comment was indeed about the content of the preprint. We used this to exclude irrelevant comments, those that merely redirected to other platforms, and those providing links to other resources with no contextualization regarding the content of the preprint. In total, we classified 192 comments (14.6% of non-authors’ comments) under this definition. Evaluators were asked to briefly describe the content of these comments, and these descriptions were used to derive categories, which are presented in **Suppl. Table 4**. Notably, 33% (n = 64) were links to other review/commenting platforms (61 of them from Oxford Immunology Network COVID-19 Literature Reviews, 1 from PreReview, 1 from Publons, and 1 from DataMethods) but the content of reviews/comments in these other platforms were not assessed. Most other comments in this category addressed the preprint’s topic or related issues but not the content of the preprint itself.

The remaining 1,125 comments were assessed using the main data collection form. As described in the Methods section, we tried to capture content elements that would be expected to be found in journal-elicited peer review, such as a summary of the manuscript, an assessment of whether conclusions are supported, compliments and criticisms. We also looked for content that would point to an interaction between readers and authors, such as asking questions, providing new data and analyses or references, and aimed to identify whether comments came from an organized review effort. **Table 1** describes these results.

**Table 1.** General content and format of comments. These categories are applicable to comments from non-authors and about the content of the preprint (1,125 comments, see Methods and Fig. 1 for details). They are not mutually exclusive; thus, the sum of percentages is greater than 100%.

| **Category** | **n** | **%**<br>**[95% C.I.]** |
| --- | --- | --- |
| Includes a criticism, correction or suggestion | 694 | **61.7**<br>[59, 65] |
| Includes a compliment or positive appraisal | 428 | **38.0**<br>[35, 41] |
| Includes a question | 393 | **34.9**<br>[32, 38] |
| Includes content that was not classified above (as criticism, correction, suggestion, compliment or question) | 69 | **6.1**<br>[5, 7] |
| Includes references | 284 | **25.2**<br>[23, 28] |
| Includes a summary description | 110 | **9.8**<br>[8, 11] |
| Explicitly questions a conclusion of the article | 98 | **8.7**<br>[7, 10] |
| Provides new data | 12 | **1.1**<br>[0, 2] |
| Provides new analyses | 5 | **0.4**<br>[0, 0.8] |
| Provides both new data and analyses | 4 | **0.4**<br>[0, 0.7] |
| Presented clearly enough for understanding | 1109 | **98.6**<br>[98, 99] |
| From an organized review effort | 75 | **6.7**<br>[5, 8] |
| Offensive (to the authors) | 2 | **0.2**<br>[0, 0.4] |

We found that 61.7% of comments (n = 694) included at least one criticism, correction or suggestion, while 38% included compliments or positive appraisals and 34.5% included questions. The overlap of these categories within comments is shown in **Suppl. Fig. 2**. Notably, a large proportion of compliments (54%) were present alongside criticisms, corrections or suggestions, but only 34% of the comments making criticisms, corrections or suggestions included a compliment.

Almost 10% of the assessed comments included a summary description of the preprint’s findings, and a similar proportion (9%) openly questioned the conclusions presented by the authors. On the other hand, very few comments included additional data or reanalysis of the data from the preprint. References were found in 25% of the assessed comments (n = 284), and most of them (65%) included journal articles, but many other reference types were identified (**Suppl. Table 5**).

Most of the comments presented their points clearly (98.6%), independently of the content itself, and only 2 comments were identified as offensive to the authors, which probably reflects moderation policies instituted by both platforms. We identified 75 comments (6.7%) as products of organized review efforts, of which 55.1% were from a single initiative (the Sinai Immunology Review Project). Other types of efforts included journal clubs, automated screening tools and graduate-level classes (**Suppl. Table 6**).

Concerning specific content, we found that criticisms, corrections or suggestions most commonly addressed the interpretation of the preprint’s results and methods, including methodological design, data collection and analysis (**Fig. 3A**). Only 1 comment was classified as a general criticism (i.e. not directed at any specific aspect of the preprint); in contrast, most compliments or positive appraisals (65.2%) were classified as general. When compliments addressed specific points of the preprint, they were mostly about the relevance and potential implications of the findings (**Fig. 3B**).

**Figure 3.**
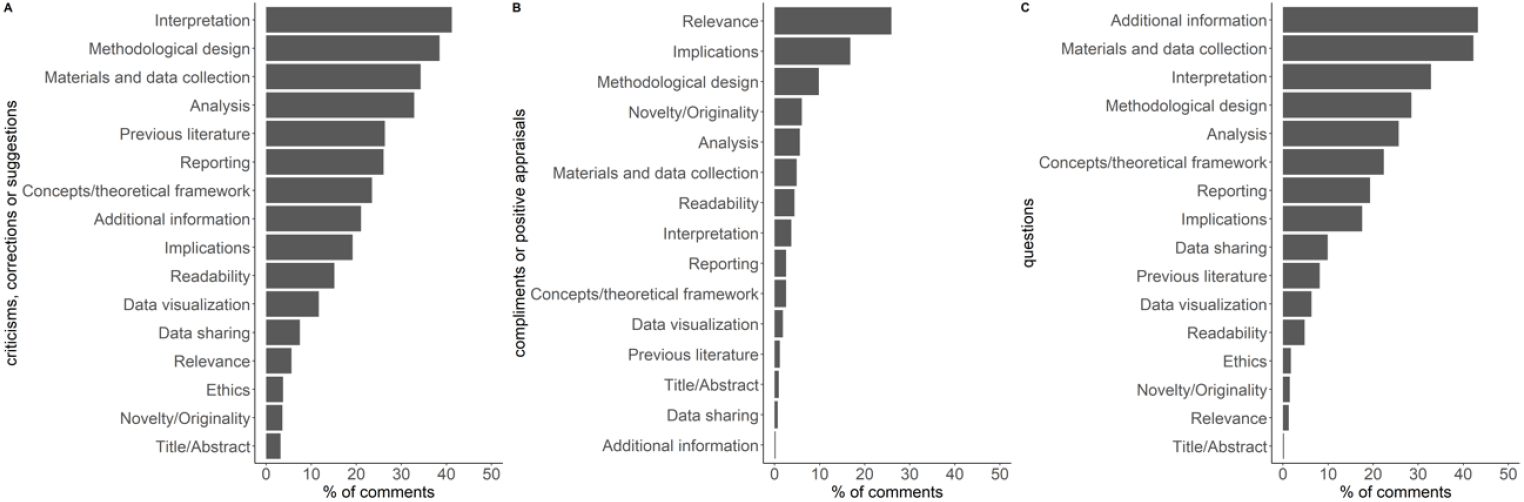
Specific content of comments. **(A)** Specific content of criticisms, corrections or suggestions. **(B)** Specific content of compliments. **(C)** Specific content of questions. Categories are not mutually exclusive (i.e. each comment could include multiple categories); thus, percentages don’t add up to 100%.

Questions mostly asked about information not present in the manuscript, which could include other results, analyses or visualizations. This was followed by questions about the materials and data collection process (**Fig. 3C**). 6.1% of comments addressed specific content without being classified as a compliment, criticism, correction, suggestion, or question. They were mostly assertions about the interpretation or implications of the preprint’s results, with no obviously positive or negative valence. **Suppl. Fig. 3** presents the aggregate results of all subcategories, and selected examples are listed in **Table 2**.

**Table 2.**
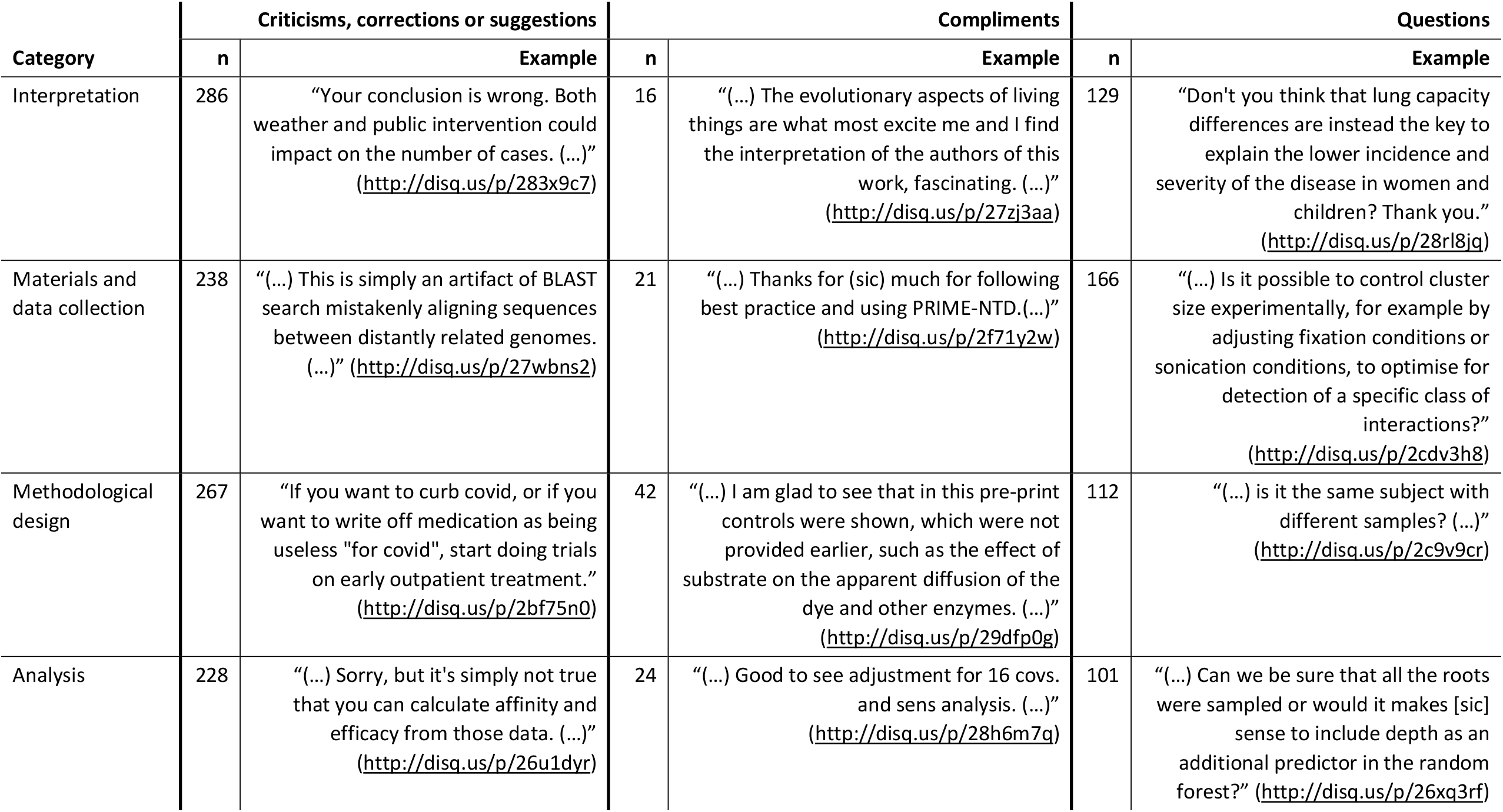
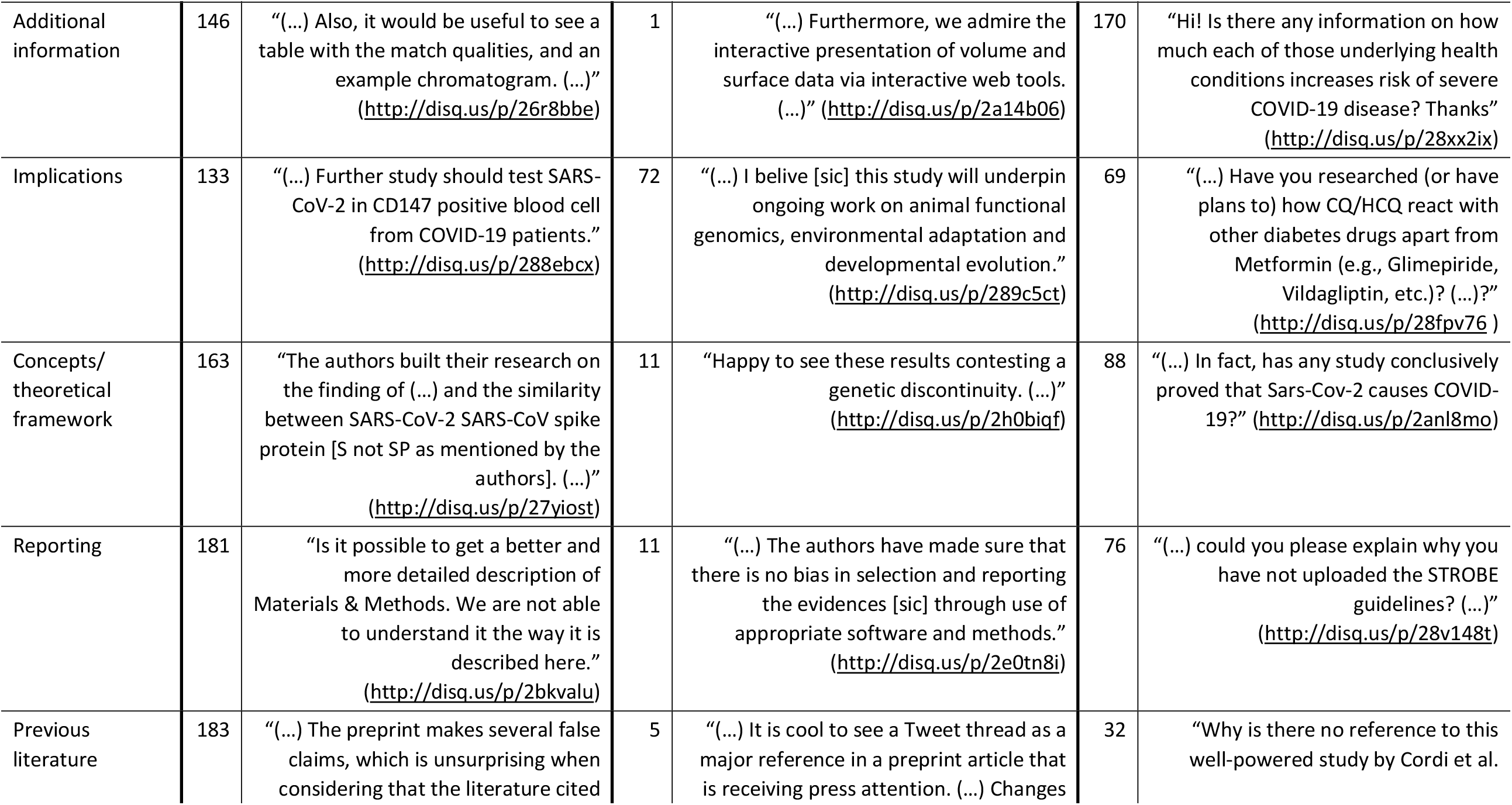
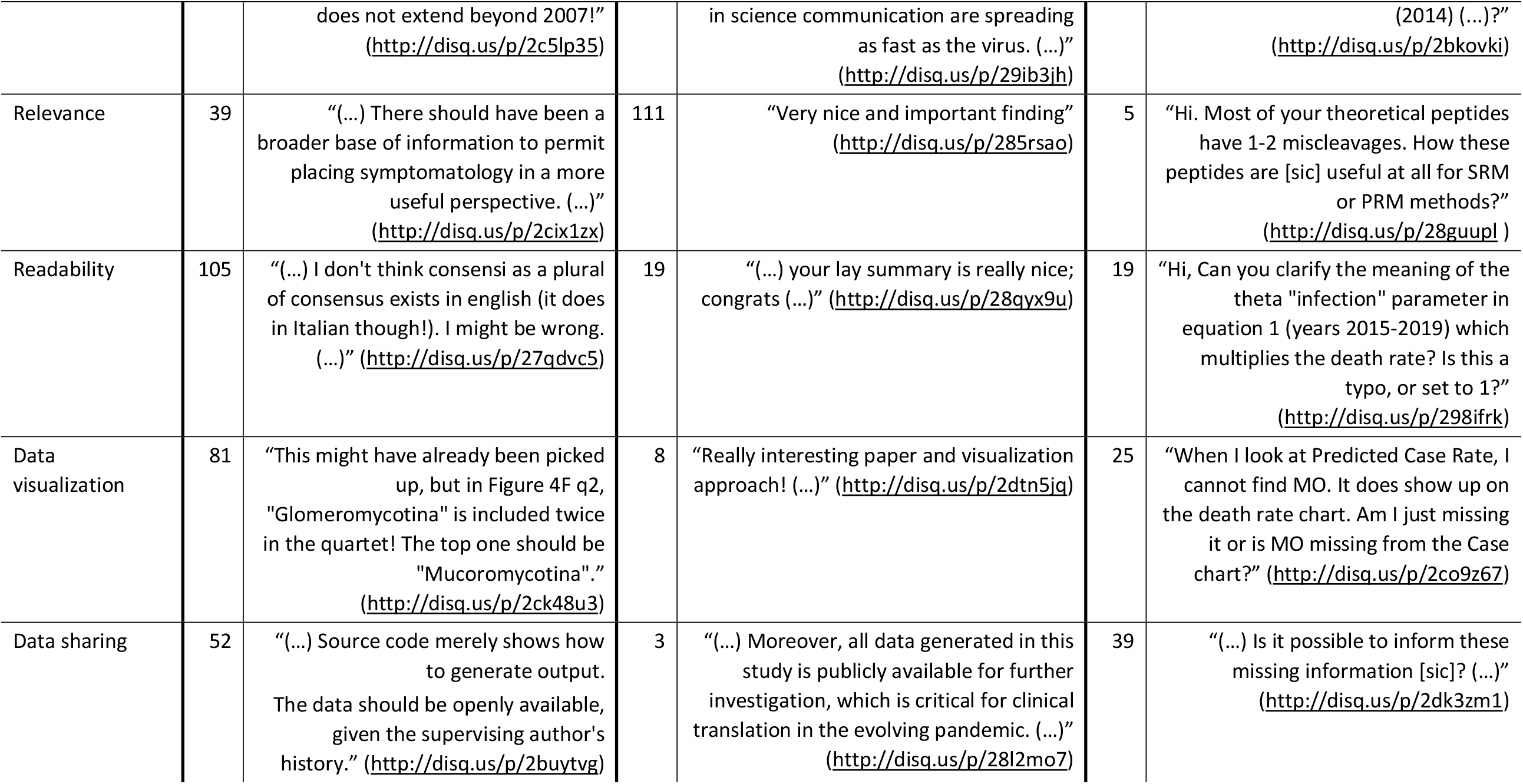
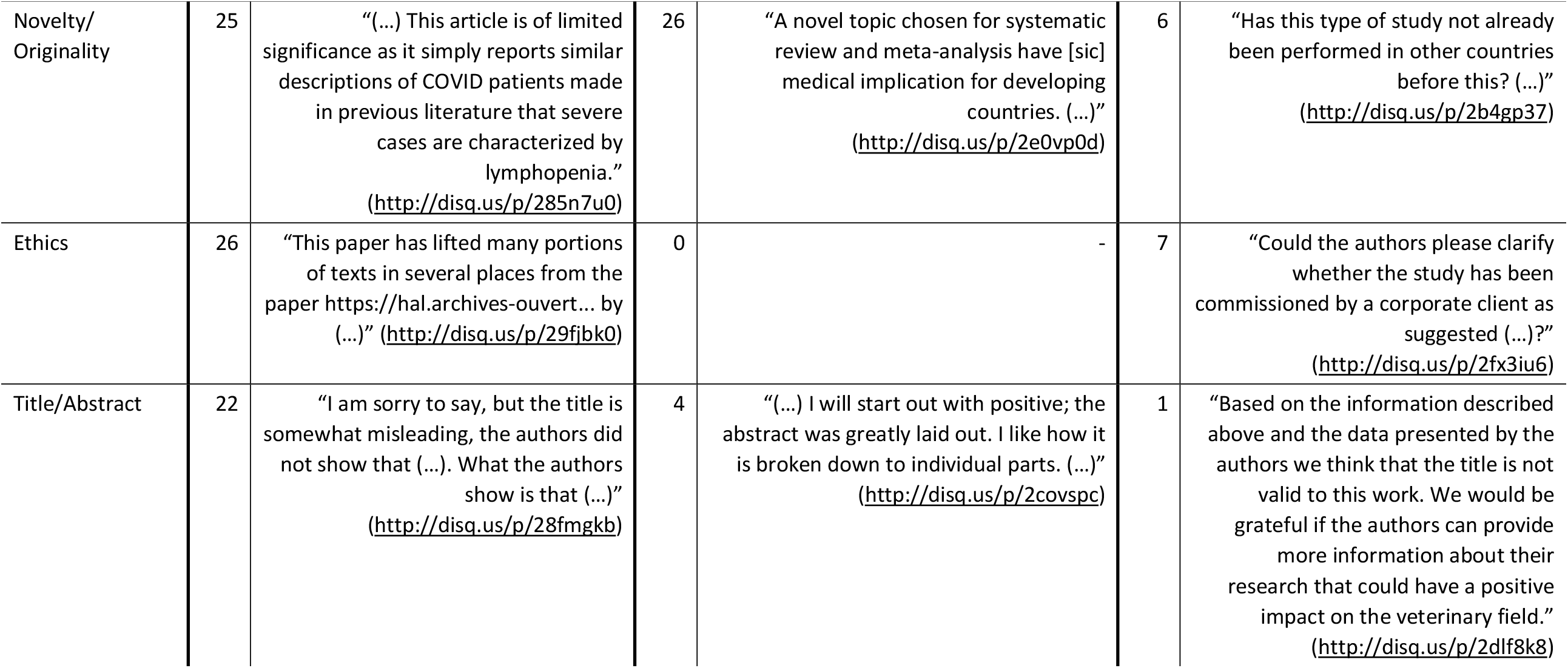
Counts and examples of specific content of comments, ordered by total number in each category. The complete text of all comments and their links are available at https://osf.io/quz6f and https://osf.io/vg6at.

General and specific content categories were also analyzed at the preprint level by aggregating all comments from each preprint (**Suppl. Table 7**). Across 810 preprints that had at least one comment from readers about their content, proportions of content types were largely similar to the analysis at the individual comment level.

### Subset analyses

During data collection, we noticed distinctive features of some types of comments that led us to analyze specific subsets (**Suppl. Table 8**). Comments from organized review efforts tended to be more similar to traditional peer review reports, and included compliments and criticisms, corrections or suggestions more often than the overall sample (52% vs. 38% and 99% vs. 62%, respectively). They also included summary descriptions and references much more often (88% vs. 10% and 44% vs. 25%, respectively).

Comments that questioned the preprint’s conclusions were predictably more critical (95% vs. 62% included criticisms, corrections or suggestions) and less positive (5.1% vs. 38% included compliments) than the overall sample. A larger proportion of these comments included references (32% vs. 25%) and provided new data, although this was still uncommon (7% vs. 1%).

We also looked at the subset of comments that elicited a response within the analyzed time frame, either from the authors or from others. These did not differ much from the complete sample, with the most pronounced change being in the proportion of comments with questions (43% among author responses and 44% among non-author responses, compared to 35% in the complete sample).

### Associations with preprint features

Finally, we studied associations between the content of comments and preprint features, presented in **Suppl. Fig. 4**. Overall, effects were small, with the most notable being that longer comments were associated with the presence of a summary description (R^2^ = 0.30, p = 1.6 × 10^−46^) and with origin from an organized review effort (R^2^ = 0.28, p = 2.5 × 10^−33^).

Longer comments were also associated with the presence of references (R^2^ = 0.06, p = 2.9 × 10^−17^) and of criticisms, corrections or suggestions (R^2^ = 0.11, p = 1.2 × 10^−35^). Preprints that had comments from authors had lower Altmetric scores and lower number of citations on average than the rest of our sample (R^2^ = 0.06, p = 2.2 × 10^−14^ and R^2^ = 0.02, p = 1.1 × 10^−5^, respectively). Preprints with COVID-related content had a smaller proportion of comments from authors (R^2^ = 0.02, p = 1.6 × 10^−6^), comments with compliments or positive appraisals (R^2^ = 0.05, p = 7.1 × 10^−18^) and comments questioning the conclusions of the study (R^2^ = 0.02, p = 1.5 × 10^−4^). They also had a higher percentage of comments that did not address the content of the preprint (R^2^ = 0.02, p = 3.6 × 10^−6^) or did not include references (R^2^ = 0.02, p = 2.1 × 10^−7^), suggesting that some of them might be outside standard academic discourse. Although these analyses were pre-planned, correlations should be interpreted as tentative due to the large number of associations in the absence of multiplicity correction.

## Discussion

In this study, we analyzed the content of 1,482 comments on 1,026 preprints on bioRxiv and medRxiv. 11% of these comments were made by authors, and these most commonly provided information about publication status. Among comments from non-authors, 85% were about the content of the preprint: of these, 62% included criticisms, corrections or suggestions, mainly about the interpretation of results and methodological aspects. Compliments were present in 38%, usually along with critical remarks and in a general form (i.e. not addressing any specific aspect of the study). Questions were present in 35% of comments and most commonly requested additional information or addressed data collection methods.

The high proportion of specific criticisms, corrections or suggestions highlight the potential of such forums to improve preprints. Similarly, questions addressing additional results or asking for clarification on methods could lead to improvements of these studies by raising attention to certain gaps. Nevertheless, we could not assess whether this potential was realized, as we did not investigate whether these comments led to changes in future versions of the preprint or journal versions.

Among comments from readers, 15% were classified as not being about the content of the preprint, including those that merely linked to another resource. About a third of those, however, were links to other peer review forums, and would in all likelihood have been classified as being about the preprint had we assessed this external content. Another third addressed the topic under study more broadly, such as questions related to health advice. For example, on a preprint about decontamination methods for N95 masks (Smith *et al*., 2020) there is a comment asking “Will an N-95 mask be safe if it is used once and then left to sit for a week then used again?” (http://disq.us/p/28o57bf). Similarly, on a preprint about the efficacy of an antiseptic preparation against the SARS-CoV-2 virus (Pelletier *et al*., 2020), a comment asks “Might this be used in conjunction with the “Naväge” device I recently saw on television that is used to clean your nasal passages?” (http://disq.us/p/29lj7×7).

This naturally brings up the question of whether the comment authors are academic researchers or members of the broader public. In many cases, comments were clearly identifiable as being from someone from the same field (e.g, http://disq.us/p/2dk3zm1) or from people outside academia (e.g., http://disq.us/p/28lpxhi). However, based on pilot assessments, we realized that making this distinction would not be possible for most comments, as the commenting service used in both preprint platforms (Disqus) allows pseudonyms and is used for many non-academic purposes as well.

Another study that analyzed comments in bioRxiv preprints (Malički *et al*., 2021) found a higher proportion of comments from authors (31%), but a largely similar distribution of content among these (i.e., focus on publication status and additional information). This study, however, looked only at preprints that had a single comment. Regarding non-author comments, a major difference between this study and ours is that it did not assess the content of comments resembling full peer review reports. This might explain the larger proportion of critical comments we found compared to that study.

A relevant question that arises is whether preprint comments could be fulfilling roles usually attributed to peer review. On average, preprint comments are certainly shorter than the average peer review report: an automated analysis of 472,449 open peer review reports from a single publisher across multiple fields of science (Buljan *et al*., 2020) reported a mean number of words of 168 for reviews recommending acceptance and 510 for those recommending major revisions in health and life sciences. Meanwhile, the mean length of more than 2.2 million open peer review reports across multiple publishers was 477 words (*Global State of Peer Review*, 2018). Although we found some large comments, the mean length in our sample of preprint comments (103 words, with a median of 43) is much smaller than these estimates.

On the other hand, comments on bioRxiv and medRxiv may be more cordial than traditional peer review. A study based on peer review reports from specific fields of biomedical research available in Publons assessed their content regarding professionalism (Gerwing *et al*., 2020). Among 920 reports, they found that 7-10% include comments demeaning or attacking the authors. In contrast, offensive comments were almost entirely absent from our sample, likely due to the moderation provided by both preprint platforms, which may be stricter than that exerted by journal editors. Notably, even the two comments classified as “offensive to the authors” in our sample (http://disq.us/p/2b1h46t and http://disq.us/p/2bf75n0) would probably not fit the definition of “offensive” in the above-cited study (Gerwing *et al*., 2020).

Concerning content, expectations about peer review vary widely (Glonti *et al*., 2019; Tennant and Ross-Hellauer, 2020). Nevertheless, various tools have been developed to assess the quality of peer review reports (Superchi *et al*., 2019), and one study identified 219 documents regarding expectations about reviewers (Glonti *et al*., 2019). According to systematic reviews of these documents, peer review reports are most often expected to assess relevance and novelty, methods (including adequacy, rigor and study design), and the interpretation of results (Glonti *et al*., 2019; Superchi *et al*., 2019). Other responsibilities include evaluation of ethical aspects, data visualization and references (Glonti *et al*., 2019).

Comments in our sample generally focused on domains similar to those described above – although this could be due to the fact that our taxonomy that was partly inspired by the above-mentioned studies. When providing criticisms, corrections or suggestions, they most often addressed interpretation, study design, data collection and analysis. Comments on relevance or novelty were mostly presented as compliments or positive appraisals. In contrast, comments on data visualization were found less frequently, and those concerning ethical aspects were very rare. It was very uncommon, however, for individual comments to address all of these aspects, and the low number of comments for most preprints suggests that this was not fulfilled by a combination of all comments either.

A limitation of our assessment is that we chose not to interpret the tone of the comments, as in pilot trials we could not reach good agreement on this topic. This led us to consider as questions anything that was explicitly phrased as a question, even if they could be interpreted as a criticism or suggestion (e.g., “(…) could you please explain why you have not uploaded the STROBE guidelines?”, http://disq.us/p/28v148t). This difficulty in reaching agreement also led us to combine criticisms, corrections and suggestions into a single category. Difficulties in distinguishing comment tone systematically are also discussed in another study on the subject (Malički *et al*., 2021).

Another limitation is that, although we built our taxonomy to describe comments that were relevant to the preprints, a non-negligible amount of them seemed to be an extension of polarized opinions on COVID-19 commonly expressed in social media. Nevertheless, many of these comments still fit our categories when taken literally. For instance, a comment questioning whether it has been proven that COVID-19 is caused by SARS-CoV-2 (http://disq.us/p/2anl8mo) was classified as a question about the theoretical framework of the preprint, despite clearly not being an academic comment.

On this topic, it is worthwhile to note that our sampling strategy deliberately excluded 25 preprints that contained more than 20 comments, most of them from medRxiv, in order not to bias our sample towards a limited number of articles. Had we included these heavily commented articles, some of which were particularly controversial (Bendavid *et al*., 2020; Magagnoli *et al*., 2020), it is likely that the proportion of non-academic discourse would have been higher.

Any discussion of the systemic role of comments as a form of post-publication peer review must take into account the fact that only 7% of the preprints in our sample of interest received any comments on the preprint servers themselves. This is less than reported by (Fraser *et al*., 2021), probably due to factors such as the time range and inclusion of non-COVID-19 preprints. Still, even their estimate for COVID-19 preprints at a time of peak attention (16%) indicates that commenting on preprints is rare – although arguably more common than post-publication comments on journals. Initiatives to recruit reviewers based on commenting and reviewing of preprints have been trying to incentivize this practice (ASAPbio, no date), but their efficacy remains an open question.

That said, it is important to realize that prevalence of feedback on preprints is likely to be higher than estimated by counting comments on the preprint pages. As of August 2022, the Reimagine Review platform listed 36 platforms and initiatives for preprint evaluation (ReimagineReview, no date). These platforms differ in who initiates the review, how reviewers are selected and whether their identities are openly available (Ettinger *et al*., 2022; ReimagineReview, no date). Some of these platforms also provide detailed guidance on how to give feedback on preprints (e.g., Hindle and Saderi, 2017; *PREreview Resource Center*, 2020), which may lead the content of comments in these platforms to be different from those on preprint platforms. Last by not least, the high Altmetric scores of some preprints suggest that much – and perhaps most – of the discussion about them happens on social media, where it can take vastly different formats, and also becomes much harder to follow.

Two collective peer review initiatives organized in response to the COVID-19 pandemic were particularly prevalent in our sample, but with an important distinction in how they provided their assessments. While the Sinai Immunology Review Project (Vabret *et al*., 2020) provided their comments directly on the commenting sections of each preprint, reports of the University of Oxford Immunology Network COVID-19 Literature Reviews project (*COVID-19 Literature Reviews — Immunology*, no date) were available in their own website, with links posted as comments to the preprint. Because of this, they were classified as “not about the content” according to our protocol and did not have their content assessed.

During our data collection period, bioRxiv and medRxiv changed the interface of their websites regarding public commenting and reviewing, with the goal of aggregating all feedback (bioRxiv, 2021). Now, both platforms include sections for community reviews (which include the Oxford Immunology Network project, Rapid Reviews: COVID-19, PreReview, PubPeer and others), transparent review led by or directed at journals (such as reviews from eLife and Review Commons, (*Transparent review in preprints*, 2019)), reports from automated screening tools (such as SciScore (Menke *et al*., 2020)) and social media mentions separated from the comment sections. Nevertheless, there is no clear distinction between content that should be posted directly as comments or via a community review platform (bioRxiv, 2022). Moreover, it is likely that much of the feedback received by preprints (such as that sent privately or articulated on social media without a direct link) remains unaggregated by these platforms, and thus disconnected from the scientific record.

In conclusion, despite being an underused mechanism for providing feedback on preprints, comments posted to bioRxiv and medRxiv have some features that resemble traditional forms of peer review. Most non-author comments are critical, and largely focused on interpretation of results and methodological aspects. References, particularly to peer-reviewed publications, are present in about a third of the assessed comments, suggesting an academic debate is taking place. Nevertheless, these comments exist for a minority of preprints, and the extent to which other post-publication review forums might be filling this gap is unclear.

Our assessment is the portrayal of a culture in transition, both in terms of adoption of preprints and of post-publication evaluations. It also describes a moment of global health emergency, leading to features that may not be maintained in the future. In the meantime, aggregating comments and reviews from multiple sources, as well as developing a structured taxonomy for classifying their content, should be regarded as high-priority goals for the improvement of scholarly communication.

## Author contributions

**Clarissa F. D. Carneiro**: Conceptualization; Methodology; Software; Formal analysis; Investigation; Writing – original draft; Visualization; Project administration. **Gabriel Costa**: Conceptualization; Methodology; Investigation; Writing – original draft. **Kleber Neves**: Conceptualization; Methodology; Software; Investigation; Writing – review & editing. **Mariana B. Abreu**: Conceptualization; Methodology; Investigation; Writing – review & editing. **Pedro B. Tan**: Software; Investigation (equal); Writing – review & editing. **Danielle Rayêe**: Investigation; Writing – review & editing. **Flávia Boos**: Investigation; Writing – review & editing. **Roberta Andrejew**: Investigation; Writing – review & editing. **Tiago Lubiana**: Investigation; Writing – review & editing. **Mario Malički**: Investigation; Writing – review & editing. **Olavo B. Amaral**: Conceptualization; Methodology; Writing – original draft; Supervision; Project administration; Funding acquisition.

