## Supplementary Tables for "Mapping the content of comments on bioRxiv and medRxiv preprints"

**Table S1** - Fleiss' kappa between the 3 evaluators for different questions in the form. Kappa values below 0.2 refer to questions with very low prevalence of one of the answers and a degree of subjectiveness, in which convergence among all 3 evaluators on infrequent answers was unlikely.

| Question | kappa | Number of comments |
|---|---|---|
| Is the comment from one of the article's authors? | 0.95 | 1922 |
| Is the (author's) comment a response? | 0.98 | 372 |
| Is the (reader's) comment a response? | 0.97 | 1503 |
| Is the comment about the content of the article? | 0.48 | 1302 |
| Is the comment from an organized review effort? | 0.84 | 960 |
| Does the comment include a summary description? | 0.72 | 955 |
| Is the comment constructed well enough? | 0.09 | 944 |
| Does the comment explicitly question any conclusion of the article? | 0.39 | 952 |
| Does the comment provide new data or analyses? | 0.15 | 957 |
| Does the comment include references? | 0.86 | 960 |
| Is the comment offensive? | 0.08 | 960 |
| Does the comment include any compliments or positive appraisals? | 0.69 | 960 |
| Does the comment include any criticisms, corrections or suggestions? | 0.63 | 959 |
| Does the comment include any questions? | 0.80 | 959 |
| Does the comment include any other specific content that was not classified as compliment, criticism, correction, suggestion or question? | 0.11 | 960 |

**Table S2** - Agreement levels across pairs of evaluators. Numbers 1 to 11 represent different evaluators. The values above the grey diagonal are the mean % agreement for all questions, while the values below the grey diagonal are the number of shared evaluations (questions) for each pair. Overall mean agreement between evaluator pairs was 92.4%.

| | **1** | **2** | **3** | **4** | **5** | **6** | **7** | **8** | **9** | **10** | **11** | **Mean** |
|---|---|---|---|---|---|---|---|---|---|---|---|---|
| **1** | | 89.7 | 90.2 | 92.3 | 91.1 | 92.2 | 92.8 | 93.3 | 92.8 | 94.7 | 91.8 | 92.1 |
| **2** | 2082 | | 89.8 | 90.4 | 91.1 | 91.1 | 91.4 | 91.8 | 88.6 | 91.3 | 89.4 | 90.5 |
| **3** | 481 | 368 | | 90.6 | 93 | 91 | 87.9 | 92 | 96 | 92.1 | 92.6 | 91.5 |
| **4** | 1934 | 1525 | 866 | | 93.7 | 94.2 | 91.8 | 94.7 | 95.7 | 92.5 | 92.5 | 92.8 |
| **5** | 2035 | 1418 | 567 | 1348 | | 93 | 90.3 | 93.1 | 90.7 | 93.5 | 93 | 92.2 |
| **6** | 1938 | 1355 | 424 | 1234 | 1322 | | 93.1 | 93.8 | 100 | 93.4 | 93.7 | 93.6 |
| **7** | 1750 | 1107 | 270 | 1107 | 891 | 1133 | | 94.5 | 91.7 | 95.1 | 92.7 | 92.1 |
| **8** | 1416 | 1062 | 578 | 1171 | 1099 | 1057 | 848 | | 92.9 | 93.7 | 93.5 | 93.3 |
| **9** | 157 | 182 | 84 | 90 | 133 | 10 | 126 | 109 | | 93.9 | 90.9 | 93.3 |
| **10** | 1220 | 886 | 594 | 1069 | 777 | 1079 | 866 | 858 | 290 | | 93.3 | 93.3 |
| **11** | 1237 | 929 | 489 | 1178 | 1135 | 1110 | 909 | 894 | 280 | 913 | | 92.3 |

**Table S3** - Specific content categories of comments posted by one of the preprint's authors. N is the number of comments including the category, and percentages refer to the 165 comments that were classified as being from an author. Categories are not mutually exclusive (i.e. each comment might include multiple themes); thus, the sum of percentages is greater than 100%.

| Category | Example | n | % |
|---|---|---|---|
| Publication status | "It has been accepted by the British Journal of Pharmacology today." (http://disq.us/p/27jtns0) | 89 | 53.9 |
| Additional information | "This is a reply to www.biorxiv.org/content/10...." (http://disq.us/p/27p3o51) | 49 | 29.7 |
| Study promotion | "(…) The tool is ideal when keeping healthcare worker safety and wellbeing perspective as priorities." (http://disq.us/p/29zk7iv) | 34 | 20.6 |
| Corrections | "There is an erratum between lines 146-149 (HLHF instead of HLLF in the stratification)." (http://disq.us/p/29gk782) | 29 | 17.6 |
| New data | "World Wildlife Fund-Mexico just announced a 2.83 ha estimate of overwinter area occupied by the eastern monarch butterfly population in Mexico this winter (…)" (http://disq.us/p/27wza05) | 17 | 10.3 |
| New analyses | "(…) We have retrained our model with confirmed cases by Feb. 11. We updated our prediction results. (…)" (http://disq.us/p/27ao9ez) | 14 | 8.5 |
| Feedback request | "(…) If you spot other mistakes, please let us know!" (http://disq.us/p/29acun8) | 12 | 7.3 |
| Extra materials | "Our new large population study is out! (…) https://www.medrxiv.org/con..." (http://disq.us/p/2cixi40) | 11 | 6.7 |

**Table S4** – Comments classified as not addressing the content of the preprint. Categories were created based on the sample, and each comment was assigned to a single category. N is the number of comments classified in each category, Percentages refer to the 192 comments that were identified as not being about the content of the preprint.

| Category | Description | Example | n | % |
|---|---|---|---|---|
| Link to external resource (peer-review) | Comments linking to other open peer-review platforms. | "I have posted a review of this on https://outbreaksci.prerevi…" (http://disq.us/p/2967lfs) | 64 | 33.3 |
| Topic under study | Comments about the topic directly under study or related topics, including health-care related questions. | "How long does it take infected individuals to develop detectable antibodies to SARS-CoV-2?" (preprint describes the development of a methodology) (http://disq.us/p/281wyp4) | 57 | 29.7 |
| Link to external resource (reference) | Comments promoting references (preprints and journal articles) without contextualization to the articles' findings. | "https://www.medrxiv.org/con...<br>Another related papers [sic] was also published recently." (http://disq.us/p/28a1cwg) | 26 | 13.5 |
| Publication status | Comments with questions or updates about peer-review or publication in a journal. | "When do you expect this to be published in a journal? I'm a little picky about formatting, so I'd love to see this work in its final form." (http://disq.us/p/2a7p3ow) | 8 | 4.2 |
| Collaboration proposal | Comments offering materials or proposing collaborations. | "If anybody wants research or visit Rohinga Camp. Please contact (…)" (http://disq.us/p/28ua732) | 7 | 3.6 |
| Scholarly communication | Comments about the use of preprints, formatting, and peer-review systems. | "I really like that you've formatted it to make it readable. It seems it would only take 10-20 minutes but it makes a huge difference. There is no real reason the preprint should be in the journal submission format." (http://disq.us/p/27jklfm) | 4 | 2.1 |
| Interaction with stakeholders (media) | Comments about how the studies were picked up by the press or social media. | "This study was cited in a nice powerpoint (…) It is being circulated on WhatsUp [sic] (…)" (http://disq.us/p/2b90iol) | 3 | 1.6 |
| Link to external resource (blog) | Comments promoting blog posts without contextualizing with the articles' findings. | "An open Letter to the editor-in-chief at JAMA on the potential efficacy of Cyclosporine in COVID-19 disease: https://medium.com/…" (http://disq.us/p/29hvf3c) | 3 | 1.6 |

| Category | Description | Example | n | % |
|---|---|---|---|---|
| Link to external resource (data) | Comments sharing databases without relating them to the article's findings. | "See https://www.ssrn.com/author… for three earlier relevant papers." (http://disq.us/p/29nliyl) | 3 | 1.6 |
| Link to external resource (media) | Comments promoting news pieces without relating them to the preprint's content. | "http://www.businessworld.in…" (http://disq.us/p/290wdok) | 3 | 1.6 |
| Access to materials | Comments about data availability without contextualizing with the preprint's content | "Is the xiFDR v2.0 software available? The Rappsilber lab link only seems to provide version 1.4.3.1" (http://disq.us/p/29iwn47) | 2 | 1.0 |
| Importance of science | Comments thanking authors for doing research. | "I'm a non health or research person. But I thank you and others who are quickly researching and reporting on observations. (…)" (http://disq.us/p/29j0m7h) | 2 | 1.0 |
| Interaction with stakeholders (policy-makers) | Comments about governmental/industry regulations related to the proposals of the study. | "Very odd FDA put this on hold" (http://disq.us/p/2bc4pil) | 2 | 1.0 |
| Public engagement | Comments asking for non-technical explanations of the study. | " I recently taught HS Biology but this is beyond my understanding. Is there anyone who can explain this study to me in simple terms?" (http://disq.us/p/28zady5) | 2 | 1.0 |
| Apology | Comment apologizing for previous criticisms. | "(…) I would like to apologize to the EV research community, the authors of this paper and specifically to (…)" (http://disq.us/p/2bpcog6) | 1 | 0.5 |
| Authorship | Questions whether a reference is by the same authors. | "https://www.preprints.org/m… Is this the same team?" (http://disq.us/p/295q4kb) | 1 | 0.5 |
| Conflict of interest | Comment about the declarations on conflicts of interest of the study. | "Note the massive conflicts of interest for these "experts." (…) Can we also get a full disclosure from Mr. Weinberger on any personal investments he may have in pharmaceutical companies or anything related to that industry." (http://disq.us/p/28y127q) | 1 | 0.5 |
| Duplication of comments | Comment about the duplication of a previous comment. | "This accidentally double-posted, due to editor approval taking longer than expected." (http://disq.us/p/2bou13j) | 1 | 0.5 |

| Category | Description | Example | n | % |
|---|---|---|---|---|
| Interaction with stakeholders (industry) | Comment about manufacturing support for the proposal of the study. | “It would be great if you would have any real manufacturing support.” (http://disq.us/p/2b60v54) | 1 | 0.5 |
| Research practices | Comment with questions and criticisms about research practices. | “These vaccine makers will take billions of dollars experimenting on us before any of their mistakes or lies can be pointed out.” (http://disq.us/p/2fkutm5) | 1 | 0.5 |

**Table S5** - Types of references cited in comments. N is the number of comments including at least one reference of each type (number of references per comment were not registered), and percentages are relative to 284 fully assessed comments with references. Comments that included isolated links with no other content were not assessed at this stage, and are described in Suppl. Table S4. Categories are not mutually exclusive (i.e. each comment could have multiple types of references); thus, percentages don't add up to 100%. References classified as 'other' included a Wikipedia page, a Google search output, a pre-registration, a presentation, an online tool and guideline documents, plus other references of unclear type.

| **Category** | **n** | **%** |
|---|---|---|
| Journal article | 186 | 65.5 |
| Preprint | 61 | 21.5 |
| Blog/website | 43 | 15.1 |
| Governmental document | 10 | 3.5 |
| Data | 8 | 2.8 |
| Manual/documentation | 8 | 2.8 |
| News piece | 5 | 1.8 |
| Social media | 3 | 1.1 |
| Book | 2 | 0.7 |
| Other | 13 | 4.6 |

**Table S6** - Types of organized review efforts. 75 assessed comments were identified as being from an organized review effort. Classifications of the type of effort were made after completion of the data collection (see Methods for details). Comments that included isolated links with no other content were not assessed at this stage, and are described in Suppl. Table S4.

| **Type of organized review effort** | **n** | **%** |
|---|---|---|
| Institutionally organized review | 55 | 73.3 |
| Lab review or journal club | 7 | 9.5 |
| Automated screening tool | 5 | 6.7 |
| Course review | 4 | 5.3 |
| Institutional journal club | 3 | 4.0 |
| Journal-requested review | 1 | 1.3 |

**Table S7** - Analysis of comment content using the preprint as the unit of analysis. N refers to the number of preprints with at least one comment falling under the category in question. % refers to the total number of preprints with at least one comment from readers that was about the content (810 preprints).

| **Category** | **n** | **%** |
|---|---|---|
| Includes a criticism, correction or suggestion | 554 | 68.4 |
| Includes a compliment or positive appraisal | 388 | 47.9 |
| Includes a question | 328 | 40.5 |
| Includes content that was not classified above | 65 | 8.0 |
| Includes references | 252 | 31.1 |
| Includes a summary description | 103 | 12.7 |
| Explicitly questions the conclusion of the article | 89 | 11.0 |
| Provides new data | 12 | 1.5 |
| Provides new analyses | 4 | 0.5 |
| Provides both new data and analyses | 4 | 0.5 |
| Presented clearly enough for understanding | 799 | 98.6 |
| From an organized review effort | 72 | 8.9 |
| Offensive (to the authors) | 2 | 0.25 |

**Table S8** – Analysis of selected subsets. The first two columns present the same data as Table 1, while all others represent subsets of this data. Percentages refer to the total number of comments in each category.

| Category | Complete dataset | | From organized review efforts (n=75) | | Questioning conclusions (n=98) | | With a response from authors (n=200) | | With a response from non-authors (n=91) | |
|---|---|---|---|---|---|---|---|---|---|---|
| | n | % | n | % | n | % | n | % | n | % |
| Includes a criticism, correction or suggestion | 694 | **61.7** | 74 | **98.7** | 93 | **94.9** | 125 | **61.6** | 56 | **61.5** |
| Includes a compliment or positive appraisal | 428 | **38.0** | 39 | **52.0** | 5 | **5.1** | 79 | **38.9** | 22 | **24.2** |
| Includes a question | 393 | **34.9** | 9 | **12.0** | 17 | **17.3** | 87 | **42.9** | 40 | **44.0** |
| Includes content that was not classified above | 69 | **6.1** | 4 | **5.3** | 0 | **0.0** | 4 | **2.0** | 6 | **6.6** |
| Includes references | 284 | **25.2** | 33 | **44.0** | 31 | **31.6** | 55 | **27.1** | 16 | **17.6** |
| Includes a summary description | 110 | **9.8** | 66 | **88.0** | 7 | **7.1** | 11 | **5.4** | 6 | **6.6** |
| Explicitly questions a conclusion of the article | 98 | **8.7** | 1 | **1.3** | - | **-** | 11 | **5.4** | 16 | **17.6** |
| Provides new data | 12 | **1.1** | 0 | **0** | 7 | **7.1** | 1 | **0.5** | 4 | **4.4** |
| Provides new analyses | 5 | **0.4** | 0 | **0** | 1 | **1.0** | 2 | **1.0** | 1 | **1.1** |
| Provides both new data and analyses | 4 | **0.4** | 0 | **0** | 2 | **2.0** | 1 | **0.5** | 0 | **0.0** |
| Presented clearly enough for understanding | 1109 | **98.6** | 75 | **100.0** | 97 | **99.0** | 200 | **98.5** | 89 | **97.8** |
| From an organized review effort | 75 | **6.7** | - | **-** | 1 | **1.0** | 5 | **2.5** | 4 | **4.4** |
| Offensive (to the authors) | 2 | **0.2** | 0 | **0** | 0 | **0** | 0 | **0** | 0 | **0** |
