## Supplementary Figures for "Mapping the content of comments on bioRxiv and medRxiv preprints"

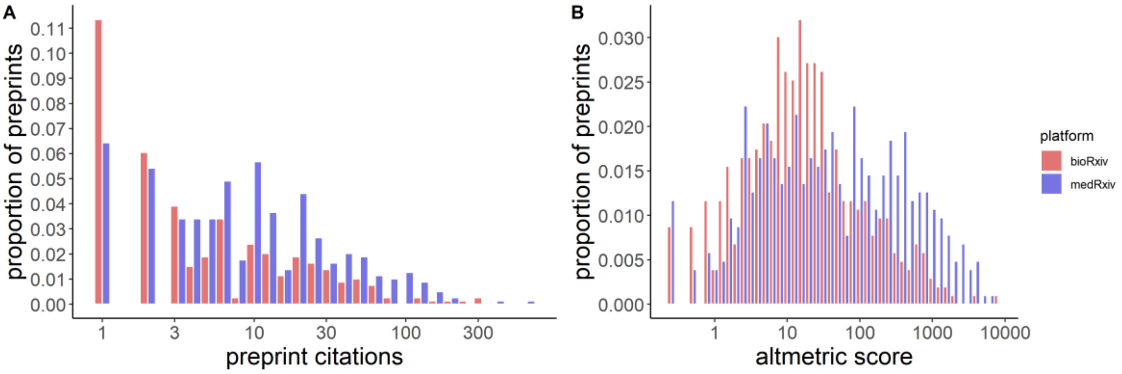


**Figure S1 – Preprint features. (A)** Distribution of number of citations to the preprint (obtained via Crossref on Oct. 28th 2021). Median (interquartile range) score is 1 (0 – 6) for bioRxiv preprints and 6 (2 – 19) for medRxiv. Overall median (interquartile range) is 3 (1 – 12). 240 preprints (23.2%) did not receive any citations. **(B)** Distribution of Altmetric scores. Median (interquartile range) score for bioRxiv preprints is 14.8 (4.6 – 48.2) and 37.0 (6.2 – 262.4) for medRxiv. Overall median (interquartile range) is 21.2 (5.5 – 119.4). Scores were unavailable for 3 preprints.

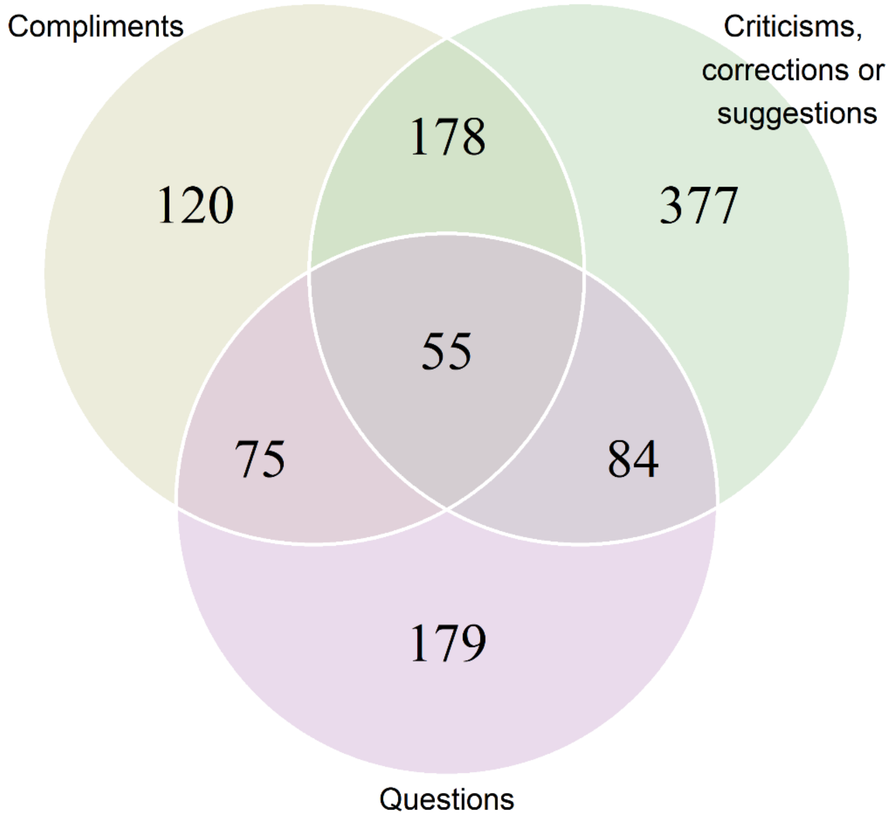


**Fig S2** - Venn diagram representing the overlap between the main content categories within comments. Note that the areas of overlap between circles are not proportional to the number of comments in each of them.

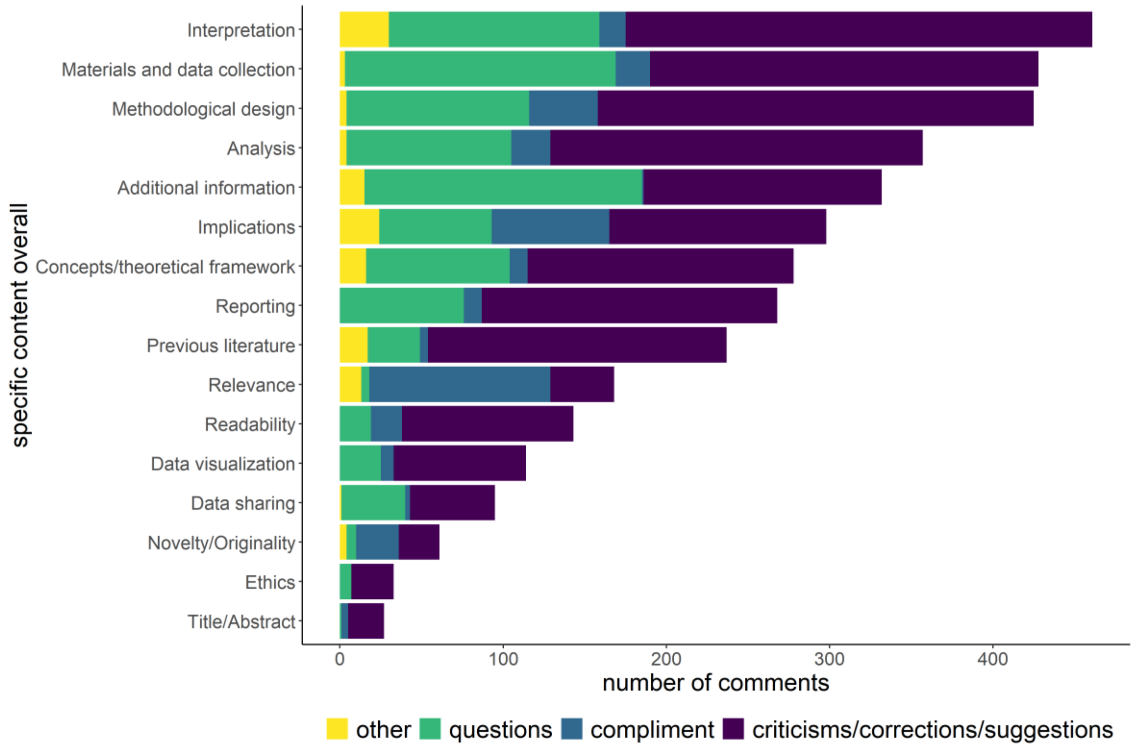


**Fig S3** – Specific content in all main categories, including criticisms/corrections/suggestions, compliments, questions and comments not classified in any of these categories (other).

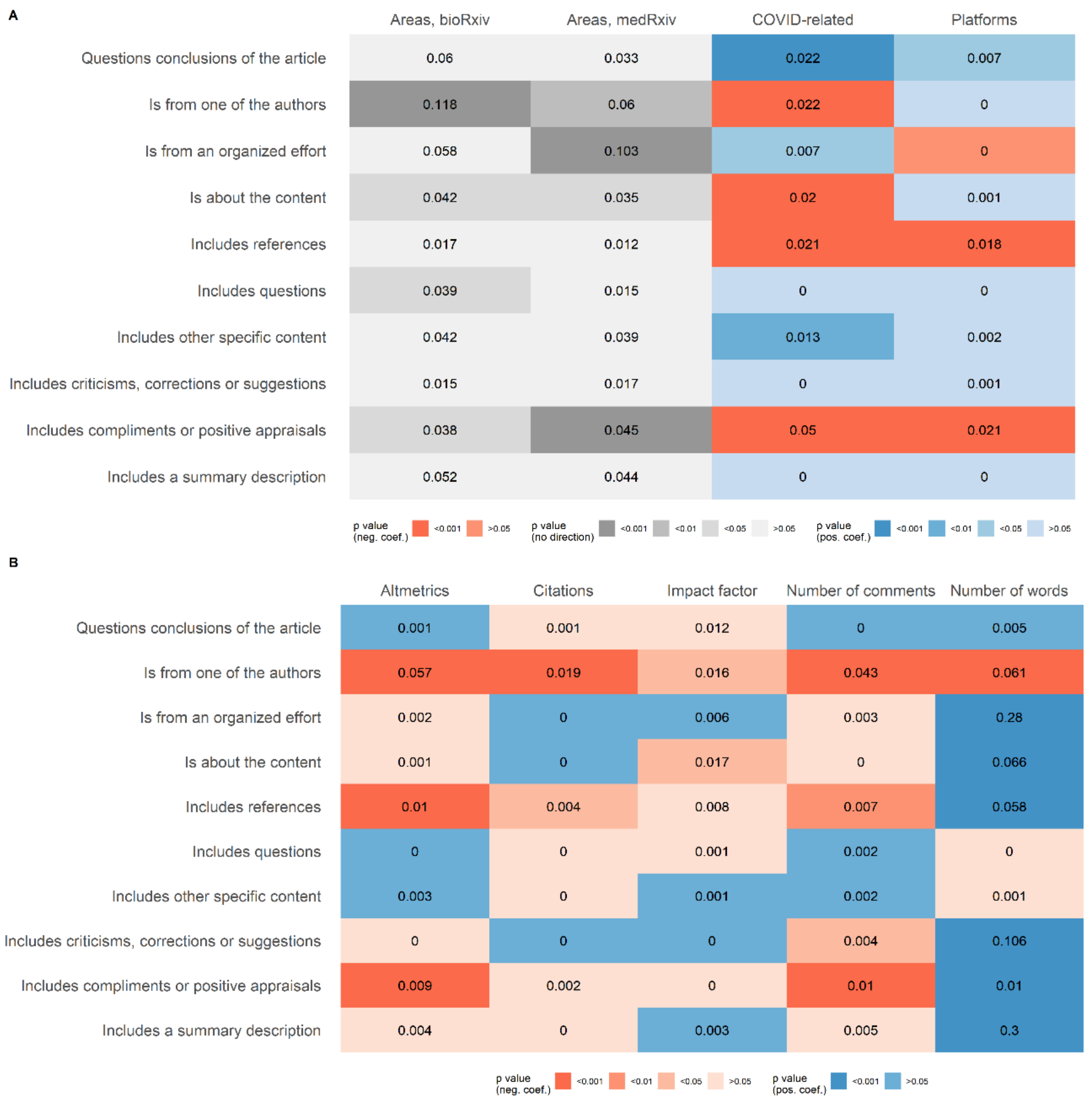


**Fig S4** - Exploratory associations between content of comments and preprint features. **(A)** Correlations with categorical variables. Tile colors indicate p value range (in red for negative associations and blue for positive associations) according to the color scale, while numbers indicate McFaden's $R^2$ values. For platforms, medRxiv is taken as reference. For COVID-related, "Yes" is taken as reference. Zero represents values smaller than 0.001. **(B)** Correlations with quantitative variables. Tile colors indicate p value range (in red for negative associations and blue for positive associations) according to the color scale, while numbers indicate McFaden's $R^2$ values. Zero represents values smaller than 0.001.
